# SELFormerMM: multimodal molecular representation learning via SELFIES, structure, text, and knowledge graph integration

**DOI:** 10.64898/2026.03.17.712340

**Authors:** Erva Ulusoy, Şevval Bostancı, Bora Engin Deniz, Tunca Doğan

**Affiliations:** Biological Data Science Lab, Dept. of Computer Engineering, Hacettepe University, Ankara, Turkey; Dept. of Bioinformatics, Graduate School of Health Sciences, Hacettepe University, Ankara, Turkey; Dept. of Health Informatics, Institute of Informatics, Hacettepe University, Ankara, Turkey

## Abstract

**Motivation:** Molecular representation learning is central to computational drug discovery. However, most existing models rely on single-modality inputs, such as molecular sequences or graphs, which capture only limited aspects of molecular behaviour. Yet unifying these modalities with complementary resources such as textual descriptions and biological interaction networks into a coherent multimodal framework remains non-trivial, hindering more informative and biologically grounded representations.

**Results:** We introduce SELFormerMM, a multimodal molecular representation learning framework that integrates SELFIES notations with structural graphs, textual descriptions, and knowledge graph– derived biological interaction data. By aligning these heterogeneous views, SELFormerMM effectively captures complementary signals that unimodal approaches often overlook. Our performance evaluation has revealed that SELFormerMM outperforms structure-, sequence-, and knowledge-based models on multiple molecular property prediction tasks. Ablation analyses further indicate that effective cross-modal alignment and modality coverage improve the model’s ability to exploit complementary information. Overall, integrating SELFIES with structural, textual, and biological context enables richer molecular representations and provides a promising framework for hypothesis-driven drug discovery.

**Availability:** SELFormerMM is available as a programmatic tool, together with datasets, pretrained models, and precomputed embeddings at https://github.com/HUBioDataLab/SELFormerMM.

## 1 Introduction

Accurate molecular representation learning lies at the core of modern computational drug discovery and cheminformatics. The ability to encode chemical compounds into informative vector representations directly impacts a wide range of downstream applications, including molecular property prediction, virtual screening, toxicity estimation, and de novo drug design. Over the past decade, advances in deep learning have significantly improved our capacity to model complex structure–property relationships, enabling data-driven approaches that complement and often surpass traditional descriptor-based methods. As molecular datasets continue to grow in scale and diversity, developing robust and transferable representation learning frameworks has become increasingly critical for reliable predictive modelling across heterogeneous biological tasks.

Recent progress in molecular representation learning has largely centred on sequence-based and graph-based approaches. Sequence-based models formulate molecular learning as a language modelling problem by treating chemical strings as textual sequences, where the derivatives of the transformer architectures is frequently employed to learn contextualised molecular representations (Fabian et al. 2020; Chithrananda et al. 2020; Ahmad et al. 2022; Irwin et al. 2022; Ross et al. 2022). The most widely used representation, SMILES (Simplified Molecular-Input Line-Entry System), encodes molecular graphs as linear character sequences (Wang et al. 2019). However, SMILES has several limitations: the same molecule can be represented by multiple non-canonical strings, syntactically valid strings may correspond to chemically invalid structures, and structural constraints are only implicitly captured (Krenn et al. 2020). To address these issues, SELFIES (Self-Referencing Embedded Strings) was introduced as a grammar-constrained representation that guarantees 100% validity and improves syntactic robustness (Krenn et al. 2020). Transformer-based models trained on SELFIES have demonstrated competitive and often superior performance to SMILES-based counterparts across various molecular property prediction tasks, highlighting the benefits of chemically constrained tokenization for molecular representation learning (Yüksel et al. 2023; Born and Manica 2023).

Molecular graph-based methods represent compounds as structured relational graphs in which atoms correspond to nodes and chemical bonds to edges. Graph neural networks operating on these molecular graphs employ message-passing mechanisms to learn embeddings that capture local chemical environments as well as higher-order topological dependencies (Yang et al. 2019; Wang et al. 2022). Variants incorporating geometric information further enhance the ability to model structure-sensitive properties (Liu et al. 2022; Zhou et al. 2023; Fang et al. 2022).

Molecular representations can also be informed by textual and structured relational knowledge sources. Text-based approaches leverage natural language descriptions, such as curated database entries or automatically generated summaries, to extract semantic embeddings using pretrained language models (Z. Liu et al. 2023; Balaji et al. 2023). Knowledge graphs represent biochemical entities, including compounds, proteins, genes, and diseases, together with their multi-relational interactions, enabling embeddings informed by relational domain knowledge (Xie et al. 2024; Doğan et al. 2021).

Although these approaches have substantially advanced molecular modelling, unimodal methods typically learn representations confined to a single molecular perspective. Sequence, structure, text, and knowledge-based encodings capture complementary aspects of molecular identity, yet they are often developed and optimised independently. Molecules are inherently multimodal entities: they possess syntactic encodings, topological structure, semantic annotations, and rich biological interaction contexts. This observation has motivated the development of multimodal learning frameworks that integrate heterogeneous molecular modalities into a unified representation space (Zhang et al. 2024; S. Liu et al. 2023; Liu et al. 2024). Nevertheless, most current approaches focus on a limited subset of modalities and primarily align views in pairwise or partially integrated settings. The incorporation of large-scale biological interaction networks and the learning of a shared embedding space that simultaneously harmonises structural, semantic, and relational information remain as promising directions for advancing multimodal molecular representation learning.

In this work, we propose SELFormerMM, a unified multimodal molecular representation learning framework that integrates SELFIES encodings, 2D molecular graphs, natural-language molecular descriptions, and biological interaction embeddings obtained from a comprehensive knowledge graph within a shared latent space. Building upon a transformer-based SELFIES encoder (Yüksel et al. 2023). SELFormerMM incorporates graph, text, and knowledge graph modalities through dedicated projection networks and aligns heterogeneous molecular views using a multi-view contrastive objective under a self-supervised learning paradigm. The model is pretrained on a large-scale multimodal dataset comprising ∼3 million molecules, with the aim of learning concise, flexible, and semantically meaningful molecular representations that generalise across diverse downstream tasks in drug discovery and development. We evaluate the quality and transferability of the learned embeddings by finetuning the model on multiple molecular property prediction benchmarks. By enabling scalable and holistic molecular representations, SELFormerMM provides a practical foundation for advancing generalizable molecular modelling strategies in computational chemistry and drug discovery.

## 2 Methods

SELFormer Multimodal (SELFormerMM) learns shared molecular embeddings by jointly leveraging sequence-based representations, structural graph information, natural language descriptions, and their multi-relational biological interactions modelled in a knowledge graph (KG) format.

SELFormerMM is trained in two stages: (i) multimodal pretraining using supervised contrastive learning and (ii) supervised finetuning for downstream molecular property prediction tasks. In the pretraining phase, the model aligns modality-specific representations of the same molecule within a shared latent space, enforcing cross-modal consistency (Fig. 1A). In the finetuning phase, the pretrained multimodal backbone is adapted to task-specific objectives by attaching a classification or regression head (Fig. 1B).

**Fig. 1.**
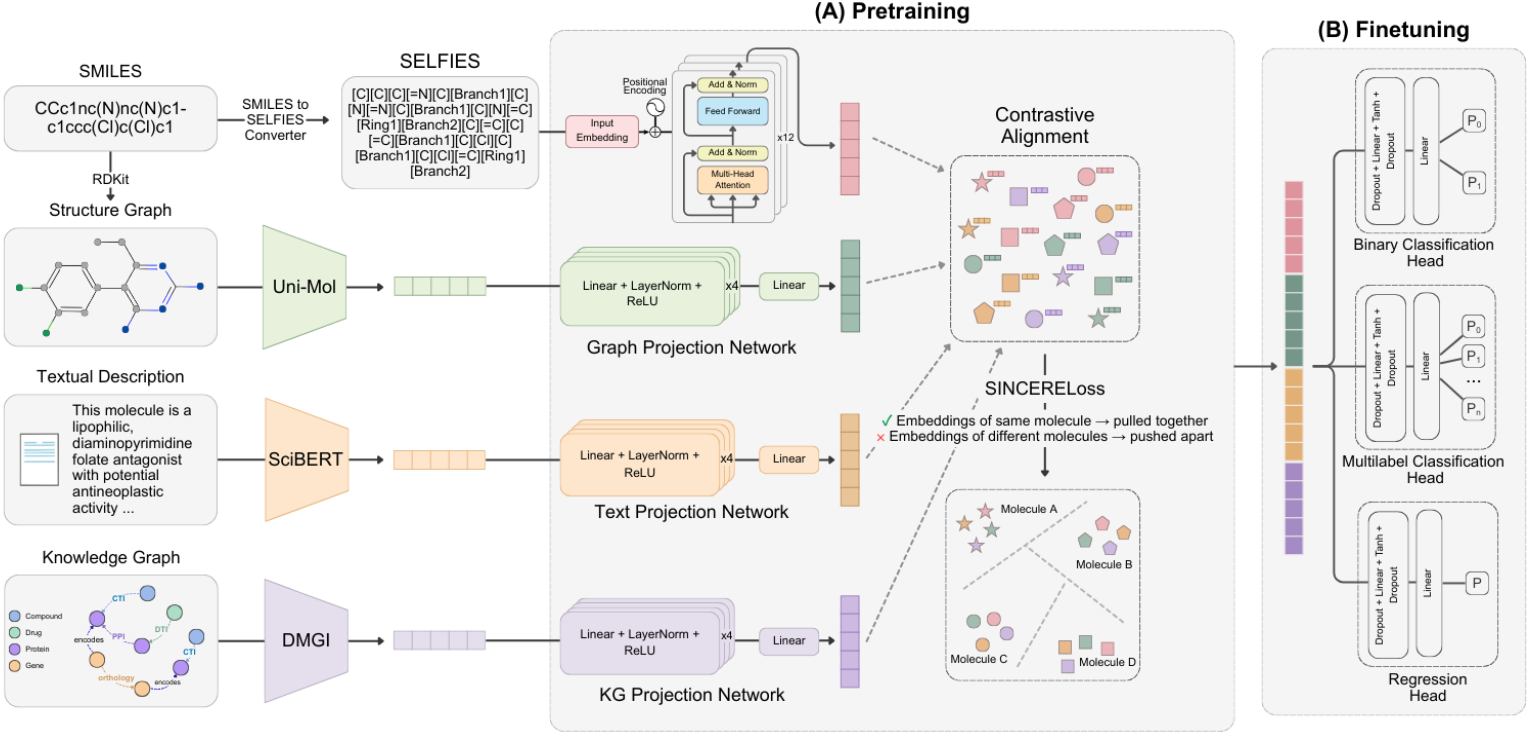
Overview of the SELFormerMM framework. **(A)** Contrastive multimodal pretraining integrates molecular information from four complementary modalities—SELFIES sequences, structural graphs, textual descriptions, and knowledge graph interactions—processed by modality-specific encoders and projection networks, then aligned in a shared representation space via contrastive learning with the SINCERE loss. **(B)** Downstream task finetuning adapts the pretrained multimodal backbone for molecular property prediction, using task-specific prediction heads for binary classification, multilabel classification, and regression.

### 2.1 Multimodal Pretraining

#### 2.1.1 Dataset

To construct the multimodal pretraining dataset, we aggregated molecular data from complementary large-scale resources, ensuring coverage of sequence, text, KG, and structure modalities.

For the molecular sequence modality, we used the ChEMBL v36 (Mendez et al. 2019) database, from which molecular SMILES representations for 2,854,815 molecules were obtained. To incorporate textual descriptions, we utilised the M3-20M dataset (Guo et al. 2024), a large-scale multimodal molecular dataset containing natural language descriptions for over 20 million molecules, derived from multiple sources including PubChem annotations, physicochemical property-based descriptions, and GPT-3.5-generated summaries. Since the molecules in M3-20M are indexed independent from ChEMBL, we constructed a mapping procedure to associate ChEMBL molecules with their corresponding textual descriptions. Mapping was first performed using standard InChIKeys, which provide a canonical chemical identifier independent of string representations. For molecules that could not be matched using InChIKeys, we applied exact SMILES matching. Remaining unmatched molecules were further processed by canonicalising SMILES strings in both datasets before attempting an additional mapping step. Through this multi-stage mapping procedure, textual descriptions were successfully assigned to 492,702 molecules.

To incorporate structured biological interaction networks of molecules, we utilised the CROssBARv2-KG (https://crossbarv2.hubiodatalab.com/, https://github.com/HUBioDataLab/CROssBARv2), which integrates large-scale biological data from 32 heterogeneous sources. To reduce memory consumption and mitigate noise during embedding generation, we constructed a focused subgraph containing only the node and edge types most directly associated with molecular properties. Specifically, we retained the node types Compound, Protein, Drug, and Gene, along with the biologically meaningful relationships connecting them: compound– target interaction (CTI), drug–target interaction (DTI), protein–protein interaction (PPI), and orthology relationships. The resulting graph contains 726,254 compound nodes, 14,594 drug nodes, 199,479 gene nodes, and 461,775 protein nodes, totalling 1,402,102 nodes. It includes 1,353,738 CTI edges, 60,500 DTI edges, 64,856 orthology edges, and 2,945,736 PPI edges, amounting to 4,424,830 relationships in total. Since CROssBARv2 represents compounds using ChEMBL identifiers, molecules in the ChEMBL v36 dataset could be directly associated with KG nodes. Through this mapping, 725,908 molecules were linked to their corresponding KG nodes.

For the structural graph modality, molecular graphs were derived directly from SMILES strings. Prior to graph construction, SMILES were validated using RDKit (Landrum 2013), and molecules for which RDKit failed to generate a valid molecule were discarded. Finally, to prepare the sequence modality for model input, all SMILES strings in the dataset were converted into SELFIES representations using the SELFIES API (Krenn et al. 2020). The distribution of modality availability across the final pretraining dataset is summarised in Table 1.

**Table 1.** Modality availability in the final pretraining dataset.

| Modality | # Molecules |
| --- | --- |
| SMILES | 2,854,815 |
| Text description | 492,702 |
| Knowledge graph | 725,908 |
| Structural graph | 2,854,734 |

#### 2.1.2 Model Architecture and Implementation

SELFormerMM comprises four modality-specific branches and a shared projection space that aligns representations from different modalities (Fig. 1A).

For the sequence modality, we employed SELFormer (Yüksel et al. 2023), a transformer-based chemical language model built upon a RoBERTa encoder implemented using the Hugging Face Transformers library. SELFIES sequences are tokenised using a bracket-delimited segmentation strategy specific to the SELFIES grammar, ensuring that each SELFIES symbol forms an atomic token. The tokenizer is implemented within a RoBERTa-compatible byte-pair encoding (BPE) framework. Special tokens follow the RoBERTa convention and are enclosed in angle brackets. As the molecule-level sequence representation, the contextualised embedding of the sequence-level special token is used. During multimodal pretraining, the SELFormer backbone and modality-specific projection networks are initialised from scratch and trained jointly, allowing the sequence encoder to adapt to the multimodal alignment objective.

For the text modality, we employed SciBERT (Beltagy et al. 2019), a pretrained natural language model based on the BERT architecture and specifically designed for scientific text. SciBERT was used to encode textual descriptions of molecules obtained from the M3-20M dataset. Description-level embeddings of dimension 768 were obtained via mean pooling over the final token-level representations of SciBERT.

For the structural graph modality, we utilised Uni-Mol (Zhou et al. 2023), a universal 3D molecular pretraining framework trained on 209 million 3D molecular conformations. Uni-Mol generates molecule-level embeddings that capture atomic connectivity and 3-dimensional structural information derived from SMILES representations. We used Uni-Mol’s molecule-level representation (cls_repr) output as the fixed-dimensional structural embedding, resulting in 512-dimensional graph representations. To encode biological interaction information, we employed the DMGI (Deep Mutual Graph Infomax) architecture (Park et al. 2020). DMGI models heterogeneous graphs using relation-specific graph convolution layers and is trained with a mutual information maximization objective that contrasts true node features with corrupted counterparts. The model was trained on the CROssBARv2-KG subgraph, with compound node features initialised using pretrained SELFormer embeddings. After training, relation-specific compound representations were aggregated via mean pooling to obtain a single 128-dimensional KG embedding per compound.

The text, structural graph, and knowledge graph encoder models were initialised from pretrained checkpoints and kept frozen during multimodal training. We defined a non-linear projection network for each of these modalities to map their native embedding spaces into the 768-dimensional hidden space of the SELFormer encoder (Fig. 1A). Each projection head is implemented as a multilayer perceptron with intermediate dimensional expansion layers, followed by LayerNorm and ReLU activations, and a final linear transformation that projects to the shared hidden dimension *H*. If a modality is unavailable for a given molecule, we feed a zero vector through the corresponding projection network, allowing the model to learn a dedicated representation for missing modalities.

To align modality-specific representations within a unified embedding space, we optimise SELFormerMM using a multi-view, multi-positive supervised contrastive objective. The objective encourages embeddings derived from different modalities of the same molecule to become similar, while pushing apart embeddings belonging to different molecules. We implement this objective using SINCERELoss, a supervised extension of the InfoNCE formulation that accommodates multiple positive views per instance (Eq. 1).

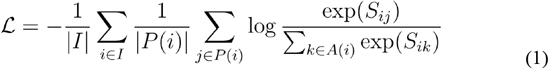

*Ei* denotes the embedding vector of the *i*-th view in the batch (e.g., sequence, graph, text, or KG representation of a molecule). *Sij* is the similarity between embeddings *E*_*i*_ and *E*_*j*_, computed as a temperature-scaled dot product with temperature parameter τ. *I* denotes the set of all anchor indices in the batch. For a given anchor *i, P(i)* is the set of positive indices corresponding to embeddings of the same molecule across different modalities, excluding the trivial self-pair. *A(i)* denotes all embeddings in the batch except the anchor itself, comprising both its positives and negatives, where negatives correspond to embeddings of different molecules.

The model was implemented in PyTorch using the Hugging Face Transformers library. SELFormerMM comprises 246,245,376 trainable parameters. Multimodal pretraining used a random 90/10 train–validation split and was optimised with the AdamW optimizer on four NVIDIA RTX A5000 GPUs. The model was trained for 25 epochs using mixed-precision optimization, requiring approximately 3 days to complete.

### 2.2 Finetuning

To evaluate the quality and transferability of the learned multimodal representations, we fine-tuned SELFormerMM on a diverse collection of molecular property prediction benchmarks from the MoleculeNet suite (Wu et al. 2018) (Fig. 1B). The selected datasets span both classification and regression tasks and cover a wide range of dataset sizes and biological endpoints. For binary classification, we used BBBP (blood–brain barrier penetration), HIV (inhibition of HIV replication), and BACE (binding to human β-secretase 1). Multilabel classification was evaluated on Tox21 (toxicity prediction) and SIDER (side effect prediction). For regression, we included ESOL (aqueous solubility), FreeSolv (hydration free energy), Lipophilicity, and PDBbind (protein–ligand binding affinity prediction). Detailed dataset statistics and modality availability for each task are provided in Table S1.

For downstream adaptation, we attach a task-specific prediction head to the pretrained SELFormerMM backbone. The backbone produces four modality-specific embeddings per molecule: the sequence representation from the RoBERTa [CLS] token and the projected embeddings of the structural graph, text, and knowledge graph modalities. These four vectors are concatenated along the feature dimension to form a single multimodal representation, which serves as input to the prediction head.

Since MoleculeNet datasets provide SMILES strings as the only molecular identifier, modality matching was performed via SMILES using the same procedure applied for the M3-20M dataset—standard InChIKey matching followed by exact SMILES matching and canonicalised SMILES matching—which nevertheless resulted in missing modalities for many samples (Table S1). Missing modalities were represented using zero vectors prior to concatenation. The prediction module maps the multimodal embedding to task-specific outputs—either class probabilities for classification tasks or a continuous value for regression tasks. To balance stability and adaptability, we adopt a partial-freezing strategy: the embedding layer and the first nine transformer blocks of the RoBERTa backbone are kept fixed, while the final three blocks, the modality projection MLPs, and the downstream head are fine-tuned. This preserves the pretrained multimodal alignment while enabling task-specific refinement at higher layers.

We follow recommended MoleculeNet protocols for evaluation. Regression and multilabel datasets are randomly split, whereas scaffold splitting is applied to the binary classification tasks using the Chemprop implementation (Heid et al. 2023). Each dataset is divided into 80/10/10 train, validation, and test splits. Hyperparameters—including backbone and head learning rates, weight decay, batch size, and number of epochs— are tuned separately for each dataset using the validation split, while final performance is reported on the held-out test split.

For classification tasks, we use cross-entropy (Eq. 2) for single label and binary cross-entropy per label for multilabel prediction.

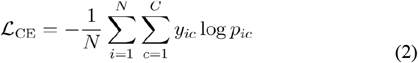

where *N* is the number of observations, *C* is the number of target classes, *y*_*ic*_ is the ground-truth indicator for sample *i* and class *c*, and *p*_*ic*_ is the predicted softmax probability for class *c*.

Model performance on classification tasks is evaluated using the Area Under the Receiver Operating Characteristic Curve (ROC-AUC). The ROC curve plots the True Positive Rate (TPR) (Eq. 3) against the False Positive Rate (FPR) (Eq. 4) across decision thresholds.

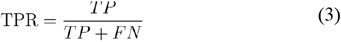

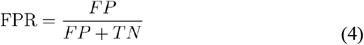

where *TP, TN, FP*, and *FN* denote true positives, true negatives, false positives, and false negatives, respectively.

For regression tasks, we optimise the Mean Squared Error (MSE) loss and report Root Mean Square Error (RMSE) for evaluation (Eq. 5).

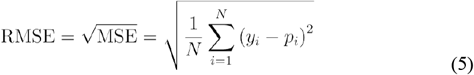

where *y*_*i*_ is the true continuous target and *p*_*i*_ is the predicted value for sample *i*. The performances are reported on the hold-out test splits.

## 3 Results

### 3.1 Pretraining and optimization

We examined how multimodal pretraining shapes the representation space on the validation split by analysing intra- and inter-molecule cosine similarities and computing their separation margin. During training, intra-molecule similarity increased substantially from 0.016 to 0.522, while inter-molecule similarity rose more moderately from 0.229 to 0.378. As a result, the separation margin improved from −0.206 to 0.144, indicating that representations of the same molecule across different modalities became more strongly aligned while preserving separation between representations of different molecules.

Local neighbourhood structure was further assessed using 5-nearest-neighbour (kNN) accuracy for molecule-identity classification in the embedding space. Validation kNN accuracy improved from 0.20 to 0.62, reflecting increasingly coherent and label-consistent clustering behaviour. Collectively, these results demonstrate that contrastive pretraining produced structured and discriminative multimodal molecular embeddings suitable for downstream adaptation.

### 3.2 Molecular property prediction benchmarks

We evaluated the effectiveness of the learned representations by fine-tuning SELFormerMM on downstream molecular property prediction tasks. We adopt the standard MoleculeNet evaluation setup (80/10/10 train/validation/test), selecting models based on validation dataset performance and reporting final results on the held-out test set.

#### 3.2.1 Classification tasks

Among the unimodal models evaluated on classification tasks (Table 2), SELFormer achieves the best performance on SIDER (ROC-AUC=0.745) and performs competitively on BBBP (0.902), closely approaching the top-performing CLM (0.915) (Born et al. 2023). This success is likely driven by the SELFIES representation, which enforces structural validity and simplifies stereochemistry, providing a robust foundation for predicting complex pharmacological responses (Yüksel et al. 2023). The strong performance of CLM, which achieves the best results on BBBP (0.915) and also performs strongly on HIV (0.813) and Tox21 (0.858), can be attributed to SMILES augmentation, which improves representation robustness and is particularly advantageous for larger datasets. On the BACE benchmark, structure-based MolCLR (0.890) (Wang et al., 2022) and the text-based MolXPT (0.884) (Z. Liu et al., 2023), benefit from large-scale pretraining on tens of millions of molecules.

**Table 2.** Performance comparison on classification-based molecular property prediction benchmarks. Results are reported as the area under the receiver operating characteristic curve (ROC-AUC) metric (higher is better). Scaffold splits are used for BACE, BBBP, and HIV, while random splits are used for Tox21 and SIDER, as suggested (Wu et al. 2018). Within each model category (unimodal and multimodal), the best-performing results are highlighted in bold, and the second-best results are indicated with underlining. “–” indicates that results were not reported for the corresponding method–task combination. Results for D-MPNN are taken from (Fang et al. 2022), while results for MGCN are taken from (Wang et al. 2022). The HIV result reported for ChemBERTa-2 is adopted from (Chithrananda et al. 2020), the previous work by the same authors, as the ChemBERTa-2 study does not report results for this dataset.

| Model type | Model name | Input type | SIDER | BACE | BBBP | HIV | Tox21 |
| --- | --- | --- | --- | --- | --- | --- | --- |
| Unimodal | MolBERT | SMILES | - | 0.866 | 0.762 | 0.783 | - |
|  | ChemBERTa-2 |  | - | 0.799 | 0.728 | 0.622 | - |
|  | CLM |  | <u>0.659</u> | 0.861 | **0.915** | **0.813** | **0.858** |
|  | D-MPNN | 2D structure | 0.646 | 0.809 | 0.710 | 0.771 | <u>0.845</u> |
|  | MolCLR |  | - | **0.890** | 0.736 | 0.806 | - |
|  | MGCN |  | - | 0.734 | 0.850 | 0.738 | - |
|  | GraphMVP-C | 3D structure | - | 0.812 | 0.724 | 0.770 | - |
|  | GEM |  | - | 0.856 | 0.724 | 0.806 | - |
|  | Uni-Mol |  | - | 0.857 | 0.729 | <u>0.808</u> | - |
|  | MolXPT | Text | - | <u>0.884</u> | 0.800 | 0.781 | - |
|  | GPT-MolBERTa |  | - | 0.734 | 0.741 | 0.755 | - |
|  | ReaKE | KG | - | 0.781 | - | 0.751 | 0.824 |
|  | SELFformer (ours) | SELFIES | **0.745** | 0.832 | <u>0.902</u> | 0.681 | 0.641 |
| Multimodal | FP-GNN | 2D structure, fingerprint | 0.661 | <u>0.860</u> | 0.916 | **0.824** | <u>0.815</u> |
|  | MoleculeSTM | 2D structure, text | - | 0.808 | **0.700** | <u>0.769</u> | - |
|  | MolLM | 2D structure, 3D structure, text | - | 0.841 | 0.757 | 0.802 | - |
|  | MvMRL | SMILES, 2D structure, fingerprint | <u>0.653</u> | **0.878** | **0.963** | - | **0.845** |
|  | GIT-Mol | SMILES, 2D structure, text | - | 0.811 | 0.739 | - | - |
|  | SELFformerMM (ours) | SELFIES, text, 2D structure, KG | **0.749** | 0.797 | <u>0.918</u> | 0.710 | 0.634 |

The competitive performance of SELFormer is further enhanced when additional modalities are aligned in SELFormerMM, particularly on the SIDER and BBBP tasks. SELFormerMM achieves the best performance on SIDER (0.749), suggesting that multimodal alignment enables the model to exploit synergistic information across modalities and better generalise biologically meaningful outcomes such as adverse-effect profiles. On BBBP, SELFormerMM achieves a score of 0.918, ranking second overall and closely following the top-performing MvMRL (0.963) (Zhang et al. 2024). MvMRL is specifically designed for molecular property prediction and incorporates SMILES, molecular graphs, and fingerprint descriptors, which encode substructural and physicochemical patterns that are particularly informative for permeability-related tasks. Despite not relying on such hand-crafted features, SELFormerMM still achieves strong performance on BBBP, indicating that it can effectively capture relevant structural signals and generalize to unseen molecular instances.

Performance on other benchmarks is more modest. On the BACE dataset, SELFormerMM achieves an ROC-AUC of 0.797 compared to 0.832 for SELFormer, which may partly relate to modality availability, as textual descriptions cover only 7% of molecules, well below the average across other benchmarks (∼65%). A similar limitation applies to the HIV dataset, where both text (14%) and knowledge graph information (0.8%) fall below the corresponding averages (∼65% and ∼6.6%). Despite this, SELFormerMM improves over SELFormer on HIV (0.710 vs. 0.681). On Tox21, SELFormerMM achieves 0.634 compared to 0.641 for SELFormer. Modelling multiple toxicity endpoints simultaneously across a large multilabel dataset may introduce additional optimization difficulty when integrating multiple modalities.

Models combining 2D molecular structures with fingerprint descriptors, such as FP-GNN (Cai et al. 2022) and MvMRL, perform strongly across tasks. Fingerprints capture predefined substructures and physicochemical patterns that complement graph connectivity, and both modalities can be directly derived from SMILES, providing broad coverage. This combination likely contributes to their stable performance across benchmarks.

#### 3.2.2 Regression tasks

For the regression benchmarks, we follow the MoleculeNet recommended random split protocol. Due to differences in the dataset splitting strategies used in prior studies, some models reported in the classification benchmarks could not be included in the regression comparison because their results are not comparable.

Among the unimodal models evaluated on regression tasks (Table 3), SELFormer achieves the best performance on ESOL (RMSE = 0.386) and comparable results on FreeSolv (1.005) and Lipophilicity (0.674). Although both SELFormer and RT (Born and Manica 2023) utilise the SELFIES representation, SELFormer consistently outperforms RT, likely due to its large-scale self-supervised pretraining followed by regression fine-tuning. On PDBbind, SELFormer (1.437) performs lower than graph-based approaches such as D-MPNN (Heid et al. 2023) and interaction-focused models, including SS-GNN (Zhang et al. 2022) and OnionNet (Zheng et al. 2019). These methods benefit from geometric representations that better capture binding affinities. Furthermore, unlike the sequence-only SELFormer, SS-GNN and OnionNet models are task-specific architectures trained directly on protein–ligand complexes.

**Table 3.** Performance comparison on regression-based molecular property prediction benchmarks. Results are reported as root mean square error (RMSE) (lower is better). Random splits are used for all datasets, following the MoleculeNet protocol (Wu et al. 2018). Within each model category (unimodal and multimodal), the best-performing results are highlighted in bold, and the second-best results are indicated with underlining. “–” indicates that results were not reported for the corresponding method–task combination. Results for D-MPNN are taken from (Fang et al. 2022).

| Model type | Model name | Input type | ESOL | FreeSolv | Lipophilicity | PDBbind |
| --- | --- | --- | --- | --- | --- | --- |
| Unimodal | MolBERT | SMILES | <u>0.531</u> | **0.941** | <u>0.561</u> | - |
|  | ChemFormer | SMILES | 0.633 | 1.230 | 0.598 | - |
|  | D-MPNN | 2D structure | 0.555 | 1.075 | **0.555** | 1.391 |
|  | SS-GNN | Protein-ligand complex | - | - | - | **1.181** |
|  | OnionNET |  | - | - | - | <u>1.278</u> |
|  | RT | SELFIES | 0.690 | 1.030 | 0.740 | - |
|  | SElFormer (ours) |  | **0.386** | <u>1.005</u> | 0.674 | 1.437 |
| Multimodal | FP-GNN | 2D structure, fingerprint | 0.675 | 0.905 | 0.625 | **1.296** |
|  | MvMRL | SMILES, 2D structure, fingerprint | **0.601** | **0.832** | 0.634 | - |
|  | SElFormerMM (ours) | SELFIES, text, 2D structure, KG | <u>0.603</u> | <u>0.883</u> | **0.574** | <u>1.328</u> |

SELFormerMM improves SELFormer’s results on PDBbind, achieving an RMSE of 1.328. This improvement may stem from the integration of both structural and knowledge graph-based information, which can provide additional biological context relevant for protein–ligand interaction tasks. On FreeSolv, SELFormerMM achieves an RMSE of 0.883, placing it behind the top-performing MvMRL (0.832).

The model also performs strongly on Lipophilicity, achieving an RMSE of 0.574, the best among multimodal models and approaching the performance of top unimodal methods. On ESOL, SELFormerMM obtains an RMSE of 0.603, remaining competitive with, or even outperforming, other multimodal frameworks. This slightly lower performance compared to the unimodal SELFormer likely reflects the challenges of multimodal optimisation. As observed in our ablation studies (Table S3), incorporating additional modalities such as text or knowledge graphs may introduce optimisation noise or gradient interference. Moreover, while multimodal alignment can benefit tasks involving complex interactions such as PDBbind, it may partially dilute the strong structural signal that the unimodal SELFormer exploits for solubility prediction.

### 3.3 Ablation study

To quantify the contribution of individual modalities and training stages, we conducted ablation experiments using identical hyperparameter configurations across all variants to ensure fair comparison. Performance was evaluated on randomly split classification and regression benchmarks.

The SELFIES-based encoder (SELFormer) served as the baseline, with additional modalities (Text, KG, Graph) incorporated sequentially via their respective projection networks. Pretraining for all model variants was limited to 20 epochs to control computational cost.

The impact of downstream finetuning was evaluated by comparing the fully multimodal pretrained model against its fine-tuned counterpart. While pretrained performance was established using a single-epoch one-pass protocol, the fine-tuned model underwent 50 epochs of optimization. Finally, we conducted leave-one-modality-out ablations to evaluate robustness under incomplete modality availability. The pretrained SELFormerMM backbone was retained, and selected modalities were masked by zeroing their embeddings during finetuning.

Across both classification (Table S2) and regression (Table S3) tasks, integrating additional modalities generally enhanced performance relative to the SELFormer baseline. An exception was observed for SIDER, where SELFormer already provided a strong baseline (ROC-AUC=0.724) and the addition of single modalities did not yield substantial improvements. However, the fully multimodal configuration achieved the best overall performance on this dataset (ROC-AUC=0.729), suggesting that integrated cross-modal representations can offer complementary benefits in certain settings. In contrast, for several other datasets, adding a single complementary modality outperformed the fully multimodal configuration. This likely reflects increased optimization difficulty during contrastive pretraining, where the alignment of multiple heterogeneous views may introduce gradient interference or cross-modal noise. Among individual modalities, 2D graph information emerged as the most robust signal, yielding the most consistent improvements and highlighting the importance of structural connectivity for molecular property prediction. This was particularly pronounced on the PDBbind dataset, where the SELFormer+Graph pretrained model (RMSE=1.333) outperformed even the fully multimodal fine-tuned variants, consistent with the strong dependence of binding affinity prediction on structural interaction patterns.

Finetuning generally improved performance across most datasets, confirming that the pretrained multimodal backbone provides a strong initialization. In contrast, performance on the Tox21 dataset remained close to random for both pretrained (ROC-AUC = 0.500) and fine-tuned variants (0.463). However, task-specific hyperparameter optimisation improves performance on this dataset (0.634; Table 2), indicating that the framework can learn meaningful signals when appropriately tuned.

Leave-one-modality-out experiments further highlight the importance of structural information. Removing the graph modality typically degraded performance, whereas removing text or knowledge graph modalities produced less consistent effects. This behaviour likely reflects differences in modality coverage: graph representations are available for nearly all molecules, while textual and knowledge graph information are available only for a subset. When a modality is available for only part of the dataset, the resulting mixture of informative and zero-valued embeddings can introduce optimization instability, bias the model toward uneven feature utilization, and create feature inconsistency across samples (Reza et al. 2025). Overall, multimodal gains depend not only on modality richness but also on coverage consistency and dataset-specific characteristics.

### 3.4 Use-case analysis

We evaluated the biochemical relevance of our molecular property predictions through case studies using representative molecules from various downstream datasets where SELFormerMM demonstrated strong performance, including BBBP, SIDER, and FreeSolv.

The BBBP task is a binary classification problem that predicts whether a compound can permeate the blood–brain barrier (BBB), a highly selective barrier that regulates the entry of molecules into the central nervous system (CNS). BBB permeability is therefore a key determinant in the development of CNS-targeting drugs. For this task, we selected two molecules indicated for neurological disorders: dextroamphetamine and benserazide. SELFormerMM predicts dextroamphetamine as BBB-permeable with a probability of 0.998. This prediction is consistent with its known pharmacological activity as a CNS stimulant used to treat narcolepsy and attention deficit hyperactivity disorder (https://go.drugbank.com/drugs/DB01576), reflecting its established ability to cross the BBB (Berman et al. 2009). In contrast, benserazide is predicted to be BBB-impermeable (0.006). Clinically, benserazide is administered with levodopa in the treatment of Parkinson’s disease and acts as a peripheral decarboxylase inhibitor that does not cross the BBB, preventing peripheral dopamine formation while allowing levodopa to reach the brain (Roche Products Limited 2020).

Case studies on the SIDER and FreeSolv benchmarks are presented in Supplementary Section 3.

## 4 Discussion

In this study, we introduced SELFormerMM, a multimodal molecular representation learning framework that integrates molecular sequences with structural graphs, textual descriptions, and knowledge graph-derived interaction information. By employing multimodal contrastive pretraining on approximately 3 million molecules, the model establishes a unified representation space that successfully integrates these heterogeneous data sources.

Evaluation on classification- and regression-based molecular property prediction benchmarks demonstrates that SELFormerMM consistently outperforms or matches state-of-the-art approaches, particularly in predicting adverse drug reactions, blood–brain barrier penetration, hydration free energy, and lipophilicity. These results suggest that integrating SELFIES with additional modalities enables the model to capture both intrinsic chemical properties and broader biological context, leading to improved predictive performance.

Ablation experiments revealed that the contribution of each modality depends strongly on its characteristics and coverage within the dataset. While structural graph information emerges as the most stable and consistently informative modality, text and knowledge graph modalities exhibit more variable effects, which can largely be attributed to their limited coverage across molecules.

Unlike prior multimodal models that either rely on fingerprints or 2D molecular graphs with near-complete coverage, or achieve high modality coverage by training on smaller curated datasets (e.g., 200–300k molecules), SELFormerMM operates at a substantially larger scale. Despite the inherent sparsity of textual and relational data, integrating these heterogeneous modalities still provides complementary signals and improves prediction performance. While datasets with complete modality coverage would enable a more seamless comparison of multimodal architectures, we use the widely adopted MoleculeNet benchmarks to ensure comparability with established baselines.

Case studies further validate that the representations learned by SELFormerMM capture biologically meaningful signals and produce predictions consistent with known pharmacological knowledge.

These findings point to several promising directions for future research. First, more robust strategies for handling missing modalities could further improve multimodal learning. Rather than relying on zero-padding, future approaches could adopt dynamic objective functions that exclude unavailable views from the contrastive loss, preventing the model from learning artificial placeholder signals. In parallel, our ongoing work explores training settings in which modality-specific encoders remain fully trainable rather than frozen, allowing end-to-end adaptation that may promote deeper cross-modal alignment. Beyond architectural improvements, improving the quality of modality-specific data may also enhance multimodal learning. Training on datasets where textual descriptions are curated from the literature may provide both better coverage and higher information quality. Extending the framework to additional downstream tasks, such as drug–target interaction prediction and de novo molecular design, represents a promising direction for demonstrating its utility in real-world drug discovery pipelines. Finally, developing a user-friendly graphical interface will be essential to making SELFormerMM accessible to experimental researchers in biology, chemistry and drug discovery.

Overall, SELFormerMM demonstrates strong potential for multimodal molecular representation learning, underscoring the need for a multifaceted approach to molecular systems. By integrating complementary chemical and biological signals, the framework produces richer and more interpretable molecular representations, enabling more effective exploration of chemical space and prioritization of biologically relevant candidates for drug discovery. To support further research, we shared SELFormerMM as a programmatic tool, together with its datasets and pretrained models, at https://github.com/HUBioDataLab/SELFormerMM.

