## Supplementary Material for "SELFormerMM: multimodal molecular representation learning via SELFIES, structure, text, and knowledge graph integration"

### 1 MoleculeNet dataset statistics

**Table S1.** Overview of the molecular property prediction tasks used for model finetuning, along with dataset statistics, modality coverage, and evaluation protocols.

| Dataset | Type of the task | # of compounds | SMILES | Text description | Knowledge graph | Structural graph | Split type | Metric |
|---|---|---|---|---|---|---|---|---|
| BBBP | Binary classification | 2039 | 2039 | 1376 | 55 | 2039 | Scaffold | ROC-AUC |
| HIV | Binary classification | 41 127 | 41 127 | 5852 | 320 | 41 120 | Scaffold | ROC-AUC |
| BACE | Binary classification | 1513 | 1513 | 112 | 198 | 1513 | Scaffold | ROC-AUC |
| SIDER | Multilabel classification | 1427 | 1427 | 1151 | 27 | 1299 | Random | ROC-AUC |
| Tox21 | Multilabel classification | 7831 | 7831 | 6011 | 361 | 7823 | Random | ROC-AUC |
| ESOL | Regression | 1128 | 1128 | 996 | 40 | 1128 | Random | RMSE |
| FreeSolv | Regression | 642 | 642 | 588 | 31 | 642 | Random | RMSE |
| Lipophilicity | Regression | 4200 | 4200 | 1352 | 1010 | 4200 | Random | RMSE |
| PDBbind | Regression | 9880 | 11 908 | 2338 | 490 | 9871 | Random | RMSE |

### 2 Ablation study results

**Table S2.** Ablation study results on classification-based molecular property prediction benchmarks. Results are reported as area under the receiver operating characteristic curve (ROC-AUC) (higher is better) using random splits for all datasets. SELFormer denotes the SELFIES-based baseline encoder. SELFormer+[Modality] indicates pretrained variants with a single additional modality integrated via modality-specific projection networks, while SELFormerMM represents the fully multimodal model. Fine-tuned variants correspond to downstream optimization of the SELFormerMM backbone, and No [Modality] denotes leave-one-modality-out settings implemented by masking the corresponding embeddings during finetuning. Best results within each model category are shown in bold.

| Model name | SIDER | BACE | BBBP | HIV | Tox21 |
|---|---|---|---|---|---|
| SELFormer (pretrained, 20 epochs) | 0.724 | 0.507 | 0.615 | 0.548 | 0.500 |
| SELFormer+Text (pretrained, 20 epochs) | 0.722 | 0.750 | 0.724 | 0.647 | 0.500 |
| SELFormer+KG (pretrained, 20 epochs) | 0.722 | 0.718 | 0.727 | 0.647 | 0.500 |
| SELFormer+Graph (pretrained, 20 epochs) | 0.724 | 0.745 | 0.748 | 0.640 | 0.500 |
| SELFormerMM (pretrained, 20 epochs) | 0.729 | 0.609 | 0.694 | 0.581 | **0.500** |
| SELFormerMM (fine-tuned) | **0.730** | 0.782 | 0.825 | 0.710 | 0.463 |
| SELFormerMM-No text (fine-tuned) | 0.696 | 0.793 | **0.845** | **0.711** | 0.463 |
| SELFormerMM-No KG (fine-tuned) | 0.696 | **0.802** | 0.801 | 0.707 | 0.463 |
| SELFormerMM-No graph (fine-tuned) | 0.701 | 0.787 | 0.818 | 0.701 | 0.463 |

**Table S3.** Ablation study results on regression-based molecular property prediction benchmarks. Performance is reported as root mean square error (RMSE) (lower is better) using random splits for all datasets. SELFormer denotes the SELFIES-based baseline encoder. SELFormer+[Modality] indicates pretrained variants with a single additional modality integrated via modality-specific projection networks, while SELFormerMM represents the fully multimodal model. Fine-tuned variants correspond to downstream optimization of the SELFormerMM backbone, and No [Modality] denotes leave-one-modality-out settings implemented by masking the corresponding embeddings during finetuning. Best results within each model category are shown in bold.

| Model name | ESOL | FreeSolv | Lipophilicity | PDBbind |
|---|---|---|---|---|
| SELFormer (pretrained, 20 epochs) | 2.812 | 5.304 | 1.208 | 1.664 |
| SELFormer+Text (pretrained, 20 epochs) | 1.927 | 4.132 | 0.938 | 1.353 |
| SELFormer+KG (pretrained, 20 epochs) | 1.880 | 4.801 | 0.991 | 1.347 |
| SELFormer+Graph (pretrained, 20 epochs) | 1.460 | 3.705 | 0.928 | **1.333** |
| SELFormerMM (pretrained, 20 epochs) | 1.468 | 3.656 | 1.015 | 1.368 |
| SELFormerMM (fine-tuned) | 0.675 | **0.927** | 0.600 | 1.341 |
| SELFormerMM-No text (fine-tuned) | **0.617** | 1.050 | **0.594** | 1.343 |
| SELFormerMM-No KG (fine-tuned) | 0.672 | 0.959 | 0.603 | 1.350 |
| SELFormerMM-No graph (fine-tuned) | 0.716 | 1.124 | 0.601 | 1.363 |

### 3 Use-case analysis: SIDER and FreeSolv benchmarks

SIDER is a multi-label classification task that groups drug side effects into 27 classes based on the affected system organ. To evaluate whether SELFormerMM captures clinically meaningful toxicity patterns, we analysed the predictions for ibandronate, a bisphosphonate commonly used to treat osteoporosis in postmenopausal women (https://go.drugbank.com/drugs/DB00710). SELFormerMM assigns high prediction scores to several clinically relevant system organ classes, including gastrointestinal disorders (0.99), musculoskeletal and connective tissue disorders (0.81), and infections and infestations (0.71). According to the SIDER database, ibandronate is associated with adverse effects such as back pain, arthralgia, dyspepsia, abdominal pain, nausea, diarrhea, and influenza-like symptoms, reported with frequencies of approximately 2–14% (http://sideeffects.embl.de/drugs/60852). These side effects map directly to the organ-system classes predicted by SELFormerMM, indicating that the model captures clinically meaningful toxicity signals. Importantly, unrelated categories such as “pregnancy, puerperium and perinatal conditions” receive low prediction scores (0.05), further indicating that the model learns biologically plausible toxicity patterns.

FreeSolv is a regression benchmark containing experimentally measured hydration free energies for small organic molecules. To assess whether SELFormerMM captures chemically meaningful solvation patterns, we analysed the agreement between predicted and experimental values across the dataset. SELFormerMM achieves a mean absolute deviation of approximately 0.21 kcal/mol, indicating close agreement with experimental measurements. For instance, the polar amine-containing molecule CCNCC (CHEMBL1189) exhibits strong hydration (experimental −4.07 kcal/mol), which the model accurately predicts (−4.07). In contrast, the hydrophobic alkyl bromide CCCCCCCCBr (CHEMBL156047) shows an unfavourable hydration free energy (0.52 kcal/mol), again closely matched by the prediction (0.53). However, fewer than 1% of the molecules in the dataset (6 out of 642) show an opposite sign between predicted and experimental hydration free energies. Most of these cases involve small halogenated compounds, such as CF and CC(F)F, which are predicted to have positive hydration free energies while experimentally exhibiting slightly negative values. Such cases likely reflect the subtle balance between hydrophobic substituents and polarization effects in aqueous environments, which can be challenging to capture consistently.
